# Adult-like thick-filament protein stoichiometry in immature human iPSC-derived cardiomyocytes

**DOI:** 10.64898/2026.11.09.761920

**Authors:** André Ventura-Gomes, Madalena Calheiros da Gama, Marta Furtado, Maria Carmo-Fonseca

## Abstract

Human induced pluripotent stem cell-derived cardiomyocytes (iPSC-CMs) retain immature sarcomeric isoforms, raising questions about their ability to recapitulate the protein stoichiometry of adult human myocardium. We integrated two independent quantitative proteomic datasets of iPSC-CMs with published fetal and region-resolved adult human heart proteomes to compare sarcomeric protein abundance. Although iPSC-CMs clustered with fetal heart based on their global shared proteome and retained fetal troponin I expression, key thick-filament protein ratios closely matched those of adult myocardium. Myosin heavy-to-light chain ratios approached the expected 2:4 stoichiometry, while myosin-binding protein abundance relative to myosin heavy chains was comparable across iPSC-CMs and adult cardiac regions. These findings reveal that developmental immaturity in sarcomeric isoform expression does not entail altered thick-filament protein proportions. Preservation of adult-like myosin-binding protein stoichiometry supports the use of iPSC-CMs to investigate sarcomeric diseases associated with altered protein dosage, particularly MYBPC3-associated hypertrophic cardiomyopathy.

---

Normal sarcomere function depends on timely developmental isoform switching and a precise stoichiometry of thin- and thick-filament components. Human induced pluripotent stem cell– derived cardiomyocytes (iPSC-CMs) are valuable cellular models for cardiovascular research. However, they retain an immature phenotype characterized by fetal sarcomeric isoforms, such as slow skeletal troponin I (TNNI1) instead of cardiac troponin I (TNNI3). Whether the relative stoichiometry of sarcomeric components in iPSC-CMs recapitulates that of the human heart remains unresolved.

To compare the proteome of *in vitro*–differentiated cardiomyocytes with that of the human heart *in vivo*, we generated two independent quantitative proteomic datasets of iPSC-CMs (Furtado et al., 2026) by liquid chromatography–tandem mass spectrometry (LC-MS/MS) using orthogonal workflows: isobaric tandem mass tag (TMT; PXD070546) multiplexing and label-free data-independent acquisition (DIA; PXD086995). These datasets were integrated with a published human fetal heart proteome (Kim et al., 2014; PXD000561) and region-resolved adult human heart proteomes from three ventricular and three atrial regions (Doll et al., 2017; PXD006675) (**Figure 1A**). Proteome depth ranged from 4478 to 9198 proteins, and 3041 proteins were quantified in all four datasets (**Figure 1B**).

**Figure 1.**
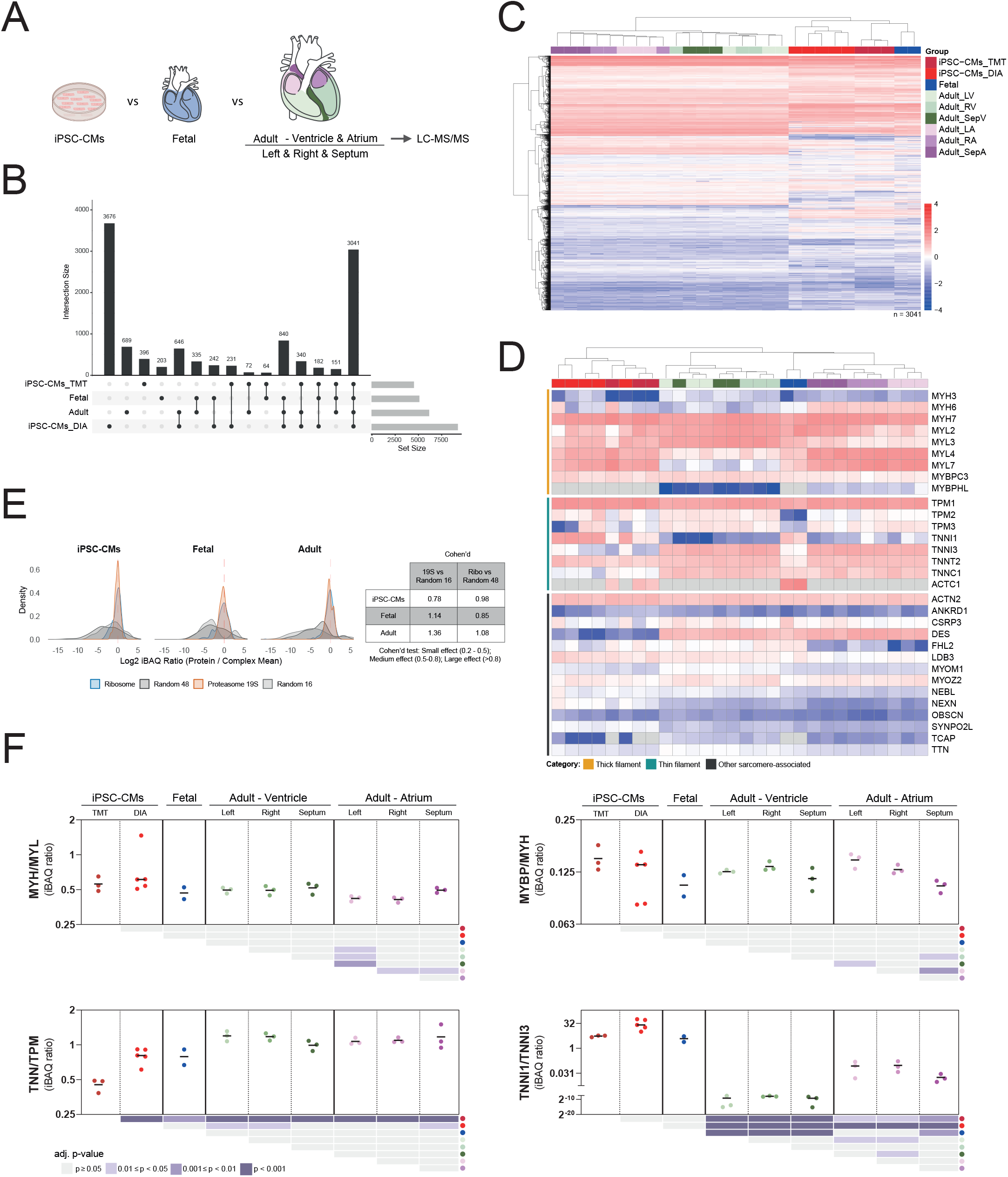
Sarcomeric stoichiometry and isoform composition in iPSC-CMs and fetal and adult human heart. **A**, Study design. iPSC-CMs were analyzed by TMT (n=3) and DIA (n=5) and compared with published proteomes of fetal heart (n=2) and 6 regions of adult heart (left and right ventricles, ventricular septum, left and right atria, and atrial septum; n=3 hearts). **B**, UpSet plot of proteins quantified per dataset. Vertical bars show intersection sizes, and horizontal bars show dataset sizes. **C**, Hierarchical clustering of 3041 proteins quantified in all datasets. **D**, Hierarchical clustering of 30 sarcomeric and sarcomere-related proteins. In C and D, iBAQ values were log2-transformed and Z-scored within each sample; grey indicates not detected or filtered out by the ≥ 2 unique peptides filter (ACTC1 shares most peptides with other actin isoforms). **E**, Density distributions of pooled log2 iBAQ ratios (protein/complex mean) for subunits of the ribosome and the 19S proteasome subunit and for size-matched random protein sets (48 and 16 proteins, respectively) in iPSC-CMs, fetal and adult heart. The dashed line marks a 1:1 ratio. Effect sizes were estimated using Cohen’s d. **F**, iBAQ ratios of troponins to tropomyosins (TNN/TPM; TNNI1/3, TNNT2, TNNC1 and TPM1, 2 and 3), myosin heavy to light chains (MYH/MYL; MYH3/6/7 and MYL2/3/4/7), TNNI1 to TNNI3, and MyBP-C and MyBP-HL to myosin heavy chains (MYBP/MYH). Points are individual samples, horizontal bars are medians, and y-axes are log2-scaled. Grids below each plot show pairwise comparisons using one-way ANOVA with Tukey-adjusted post hoc tests for independent groups. Comparisons between paired adult samples were analysed using a linear model that blocks for subject, followed by Tukey-adjusted post hoc tests.

Unsupervised hierarchical clustering of Z-scored intensity-based absolute quantification (iBAQ) values for the 3041 shared proteins separated adult samples from iPSC-CMs and fetal heart (**Figure 1C**). Adult samples segregated into atrial and ventricular subclusters, whereas iPSC-CMs grouped with fetal heart. Restricting the analysis to 30 sarcomeric and sarcomere-related proteins separated iPSC-CMs from all tissue samples and grouped fetal heart with adult atria, consistent with fetal expression of atrial-type myosin light chains (MYL4, MYL7). iPSC-CMs expressed lower levels of TNNI3, telethonin (TCAP) and desmin (**Figure 1D**).

To test whether iBAQ ratios recover known stoichiometries, we examined subunits of two stable equimolar complexes, the ribosome and the 19S proteasome (**Figure 1E**). In all three groups, subunit iBAQ ratios formed narrow distributions centered at log2 = 0 (1:1), in contrast to the broader distributions of size-matched random protein sets (Cohen’s d, 0.78–1.36).

We compared three stoichiometric ratios and one isoform ratio across the 9 sample groups (**Figure 1F**). Myosin heavy and light chains assemble at a fixed 1:2 molar ratio (two heavy and four light chains per myosin hexamer). MYH/MYL ratios approached the expected 0.5 in both iPSC-CMs datasets (mean of both groups: ∼0.57), fetal (∼0.47), and adult ventricles (∼0.50) and atria (∼0.47), with no difference between iPSC-CMs and any adult region (adj. p-value ≥ 0.05). The troponin-to-tropomyosin ratio (TNN/TPM) was stable across adult regions (∼1.0–1.2) but systematically below the 1.5 predicted by one troponin heterotrimer per tropomyosin dimer. In iPSC-CMs, TNN/TPM approached fetal values in the DIA dataset (∼0.8) but was reduced in the TMT dataset (∼0.4; adj. p-value <0.01 vs all other groups). Atria co-express MyBP-H-like (MYBPHL), which occupies the same C-zone sites as cardiac MyBP-C (Barefield et al., 2023); MYBPHL was not detected in iPSC-CMs or fetal heart. Total MyBP per myosin heavy chain, (MYBPC3 + MYBPHL)/MYH, was comparable in adult ventricles (0.129), atria (0.127) and iPSC-CMs (TMT, 0.141; DIA, 0.138), with no difference between iPSC-CMs or fetal heart (0.105) and any adult region (adj. p-value ≥0.05). In contrast, TNNI1/TNNI3 in iPSC-CMs did not differ from fetal values (adj. p-value ≥0.05) and exceeded that of adult ventricles by more than four orders of magnitude (adj. p-value <0.001). The human cardiac thick-filament structure places 27 MyBP-C molecules per 294 myosin heavy chains in each half filament (Dutta et al., 2023), predicting a ratio of ∼0.09. Adult tissue and iPSC-CMs exceeded this value by ∼40–50%, potentially reflecting protein-specific iBAQ quantification biases, or suggesting that a fraction of MyBP-C participates in molecular assemblies beyond its canonical association with thick filaments.

Together, these findings reveal that, despite retaining a fetal-like thin-filament isoform profile, iPSC-CMs establish near-adult stoichiometric proportions of key thick-filament components. In particular, the abundance of myosin-binding proteins relative to myosin heavy chains closely matches that observed in adult human myocardium, indicating that developmental immaturity does not necessarily entail altered sarcomeric protein stoichiometry. This finding has important implications for modelling hypertrophic cardiomyopathy, in which pathogenic MYBPC3 variants frequently cause haploinsufficiency (Marston et al., 2009). By establishing that control iPSC-CMs maintain physiological MyBP-to-myosin proportions, our results support their suitability for investigating disease-associated deficits in cMyBP-C abundance and their consequences for sarcomere function.

Limitations of this study include small sample sizes, differences in proteomic acquisition platforms and depth of coverage, and contributions from non-myocytes in cardiac tissue samples. Because iBAQ depends on peptide detectability, and homologous isoforms are quantified using different unique peptides, ratios involving isoform mixtures may vary with isoform composition. Thus, iBAQ-derived ratios approximate relative molar abundance rather than absolute stoichiometry. Nevertheless, our findings demonstrate that iPSC-CMs preserve adult-like thick-filament protein proportions despite their developmental immaturity, supporting their use as models of sarcomeric disease, particularly MYBPC3-associated hypertrophic cardiomyopathy.

## Materials & Methods

### Sample preparation for TMT hiPSCs-CM samples

Cardiomyocytes differentiated from hiPSC lines D, T and C were collected at day 30 of differentiation. The resulting pellet was stored at −80 °C and sent for sample preparation and LC-MS3 analysis to the VIB Proteomics Core in Belgium. Proteins were reduced, alkylated and digested with trypsin (1/50, w/w) overnight at 37°C. The resulting peptides were labelled for 1 hour at RT using 0.25 mg of TMTProTM labels (labels used: 127N - hiPSC line C, 128C - hiPSC line T and 133N - hiPSC line D). Fractions were collected every minute between 20 and 92 minutes and pooled every 3 minutes to generate a total of 24 samples for LC-MS3 analysis. Purified peptides were injected on an Ultimate 3000 RSLCnano system in-line connected to an Orbitrap Fusion Lumos mass spectrometer equipped with a pneu-Nimbus dual ion source

(Phoenix S&T). The mass spectrometer was operated in data-dependent mode (DDA) with a top speed of three seconds. Full-scan MS spectra (375-1500 m/z) were acquired at a resolution of 120,000 in the Orbitrap analyzer. The precursor ions were selected in the quadrupole with an isolation window of 0.7 Da and accumulated to an AGC target of 1E4 or a maximum injection time of 50 ms and activated using CID fragmentation (35% NCE). The 10 most intense MS2 fragments were selected in the quadrupole using MS3 multi-notch isolation windows of 2 m/z. The mass spectrometry proteomics raw data files have been deposited in the ProteomeXchange Consortium via the PRIDE partner repository with the dataset identifier PXD070546.

### Sample preparation for label-free DIA hiPSCs-CM samples

Cell pellets were lysed in 75 µl of 5% SDS in 50 mM triethylammonium bicarbonate (TEAB, pH 8.5). Lysates were transferred to a PIXUL plate (Active Motif) in portions of no more than 100 µl and sonicated for 5 min with the PIXUL Multi-Sample Sonicator (N = 50 pulses, pulse repetition frequency 1 kHz, burst rate 20 Hz). Plates were centrifuged at maximum speed, and cleared supernatants were transferred to a Protein LoBind plate (Eppendorf). Protein concentration was estimated with the BCA assay (Thermo Scientific). Proteins were reduced and alkylated with 10 mM TCEP and 40 mM chloroacetamide (CAA) for 30 min at 25 °C in the dark with shaking at 750 rpm. Samples were acidified with phosphoric acid to a final concentration of 2.75%. Then 495 µl of S-Trap binding buffer (90% methanol, 100 mM TEAB, pH 7.55) was added. Approximately two-thirds of each sample (∼50 µg protein) was loaded onto S-Trap micro columns (ProtiFi) in 200 µl portions, with centrifugation for 30 s at 4,000 ×g after each portion. Columns were washed three times with 150 µl of binding buffer (2 min at 1,500 ×g), rotating the column 180° before washing, and then centrifuged for an additional 1 min at 4,000 ×g. Proteins were digested on-column overnight at 37 °C with 1 µg of trypsin in 20 µl of 50 mM TEAB. Peptides were eluted sequentially with 40 µl of 50 mM TEAB, 40 µl of 0.2% formic acid, and 40 µl of 50% acetonitrile with 0.2% formic acid, each by centrifugation for 1 min at 4,000 ×g. Eluates were pooled, dried in a vacuum centrifuge and stored at −20 °C until LC-MS/MS analysis.

Peptides were re-dissolved in 20 µl loading solvent A (0.1% trifluoroacetic acid in water/acetonitrile (ACN) (99.5:0.5, v/v)) of which 2 µl was injected for LC-MS/MS analysis on an Ultimate 3000 ProFlow nanoLC system in-line connected to a Orbitrap Exploris 240 mass spectrometer (Thermo). Trapping was performed at 20 μl/min for 2 min in loading solvent A on a 5 mm trapping column (Thermo scientific, 300 μm internal diameter (I.D.), 5 μm beads). The peptides were separated on an in-house produced column (75 µm x 250 mm), equipped with a laser pulled electrospray tip using a P-2000 Laser Based Micropipette Puller (Sutter Instruments), packed in-needle with ReproSil-Pur basic 1.5 µm silica particles (Dr. Maisch) kept at a constant temperature of 45°C. Peptides were eluted by a gradient reaching 26.4% MS solvent B (0.1% FA in acetonitrile) after 80 min, 44% MS solvent B at 95 min, 56% MS solvent B at 100 min followed by a 5-minutes wash at 56% MS solvent B and re-equilibration with MS solvent A (0.1% FA in water). The flow rate was set to 250 nl/min. The mass spectrometer was operated in data-independent mode, automatically switching between MS and MS/MS acquisition. Full-scan MS spectra ranging from 375-1500 m/z with a normalized target value of 300%, a maximum fill time of 25 ms and a resolution of 60,000 were followed by 30 quadrupole isolations with a precursor isolation width of 10 m/z for HCD fragmentation at an NCE of 30% after filling the trap at a normalized target value of 2000% for maximum injection time of 45 ms. MS2 spectra were acquired at a resolution of 15,000 with a scan range of 200-1800 m/z in the Orbitrap analyser without multiplexing. The isolation intervals were set from 400 – 700 m/z with a width of 10 m/z using window placement optimization. The polydimethylcyclosiloxane background ion at 445.120028 Da was used for internal calibration (lock mass). QCloud was used to control instrument longitudinal performance. The mass spectrometry proteomics raw data files have been deposited in the ProteomeXchange Consortium via the PRIDE partner repository with the dataset identifier PXD086995.

### Raw files processing for protein group identification and IBAQ quantification

Protein group identification of all raw files was performed using MaxQuant software (version 2.6.4.0) with mainly default search settings for the different types of files, including a false discovery rate set at 1% on the peptide and protein level. All independent searches were performed against the human UniProt canonical database (UP000005640_9606; release version 2025_04). Enzyme specificity was set to cleavage C-terminal to arginine and lysine residues, including cleavage at Arg/Lys–Pro bonds, allowing up to two missed cleavages. Carbamidomethylation of cysteine was set as a fixed modification, while methionine oxidation and N-terminal acetylation were included as variable modifications. The “match between runs” feature was enabled to enhance peptide identification across LC-MS/MS runs. Intensity-based absolute quantification (iBAQ) was enabled to estimate relative molar protein abundance. Data filtering was performed using Perseus (v2.1.2.0). After log2 IBAQ transformation, proteins were filtered to require at least 3 valid values across different replicates and at least 2 unique peptides per protein group. Missing values were replaced by sampling from a normal distribution with a down shift of 1.8 standard deviations and a width of 0.3 (relative to the standard deviation of the measured values).

